# Body mass and growth rates predict protein intake across animals

**DOI:** 10.1101/2023.06.20.545784

**Authors:** Stav Talal, Jon F. Harrison, Ruth Farington, Jacob P. Youngblood, Hector E. Medina, Rick Overson, Arianne J. Cease

## Abstract

Organisms require dietary macronutrients in specific ratios to maximize performance, and variation in macronutrient requirements plays a central role in niche determination. Although it is well-recognized that development and body size can have strong and predictable effects on many aspects of organismal function, we lack a predictive understanding of ontogenetic or scaling effects on macronutrient intake. We determined protein and carbohydrate intake throughout development on lab populations of locusts and tested whether lab responses can predict results for field populations. Self-selected protein:carbohydrate targets declined dramatically through ontogeny, due primarily to declines in mass-specific protein consumption rates which were highly correlated with declines in specific growth rates. Importantly, lab results for protein consumption rates predicted results for field-collected locusts. However, field locusts consumed nearly double the carbohydrate, likely due to higher activity and metabolic rates. Combining our results with the available data for animals, both across species and during ontogeny, protein consumption scaled predictably and hypometrically, demonstrating a new scaling rule key for understanding nutritional ecology.

## Introduction

Every animal must acquire a proper balance of macronutrients to maximize their performance (Simpson and Raubenheimer, 2012). For all animals, protein is the main building block for growing tissues, and lipids and carbohydrates (non-protein) are the primary energy fuels. Comparative studies show that different animal species have a wide range of unique protein:carbohydrate (and/or lipid) targets that optimize growth, survival, and reproduction, and these are often thought of as species-specific (Behmer and Joern, 2008; Behmer, 2009; Hewson-Hughes et al., 2013). While a few studies indicate developmental effects on macronutrient intake, we lack a clear understanding about how and why ontogeny or body size affect macronutrient consumption and intake targets (Peters, 1983). To address this lack, we determined the effect of ontogeny and body mass on macronutrient (protein and carbohydrate) intake and growth rate for the polyphagous and transboundary migratory pest, *Schistocerca cancellata* (Serville, 1838), the South American locust, and integrated our results with prior studies of this topic in animals.

Foraging decisions are often driven by the need to balance protein (p) with non-protein (np) energy because these macronutrients make up the vast majority of a consumer’s diet and food sources rarely match the balance needed. The relative macronutrient requirements of individuals across development and the factors that influence these intake targets have profound implications for population dynamics and ecosystems, particularly for herbivores. For example, in many cases, growth and population levels of freshwater invertebrate herbivores are limited by protein (or more specifically, essential amino acid) availability (Fink et al., 2011). In contrast, late-instars of grasshoppers and some lepidopteran caterpillars have low protein to carbohydrate targets due to their high energy requirements for adult flight (Lee et al., 2004; Talal et al., 2020). In these cases, low nitrogen environments which harbor low-protein, high-carbohydrate plants promote outbreaks and devastating locust migratory swarms (Cease et al., 2012). Animals restricted to feeding on foods that diverge from their required p:np balance can experience pronounced performance deficits in development time, mass, reproduction and survival (Behmer and Joern, 2008; Behmer, 2009; Raubenheimer et al., 2022; Simpson and Raubenheimer, 2012; Talal et al., 2020).

The Geometric Framework for Nutrition (Simpson and Raubenheimer, 2012) was developed to study how organisms balance multiple nutrients, and identifying intake targets is a key principle. Most organisms will self-select a balance of nutrients, and this can be tested by giving individuals a choice between two or more diets differing in the ratio of two or more nutrients. Individuals differentially eat the diets to achieve an intake target. Macronutrient targets can vary across species. For example, cats selectively consume and perform better on more protein-biased diets (52p:48np) than dogs, for which a 30p:70np diet is optimal (Hewson-Hughes et al., 2013, 2011). Such variation in nutritional targets can occur even among closely related species. For example, late-instar juveniles of seven species of congeneric grasshoppers that share the same habitat exhibit widely different species-specific p:np targets that maximize their growth performance (Behmer and Joern, 2008). However, the ultimate and proximate explanations of the wide interspecific variation in macronutrient consumption remain poorly understood.

Some evidence suggests that intake of protein relative to carbohydrate (and or lipids) may generally decrease through ontogeny. Stable isotope analysis of tooth and skin suggested that mass-specific protein consumption declines during ontogeny in bottlenose dolphins (*Tursiops truncates,* (Knoff et al., 2008)). Similarly, in turtles (*Trachemys scripta*, (Bouchard and Bjorndal, 2006)), lizards (*Stellagama stellio*, (Karameta et al., 2017)), and sturgeons (*Acipenser persicu*s, (Babaei et al., 2011)), food preference, digestive efficiency, and digestive enzymatic activities indicate decreasing mass-specific protein assimilation and need as ontogeny progresses. Changes in milk composition through ontogeny also suggest that offspring nutritional requirements shift from protein-biased to energy (lipid + carbohydrate)-biased with age. Milk p:np decreases as offspring develop by various methods in different species. In northern elephant seals (*Mirounga angustirostris*), the lipid concentration of milk increases by ∼5 fold during 30 days of lactation (Riedman and Ortiz, 1979), while in humans, the protein concentration of milk decreases as lactation progresses (Ballard and Morrow, 2013; Ghabrial et al., 2011). Tammar wallabies provide milk with a lower p:np ratio to older offspring, even when nursing two offspring simultaneously (Nicholas et al., 1997). A few studies have tested for an effect of ontogeny on preferred p:np consumption and utilization in invertebrates, but generally only over a short span of the life cycle. A study of brown-banded cockroaches showed that the self-selected ratio of casein:glucose decreased from third to final instar (Cohen et al., 1987). Lepidopteran caterpillars decrease p:np consumption over three instars (Stockhoff, 1993). We lack whole-ontogeny studies of macronutrient targets, and an understanding of whether a decline in mass-specific protein intake is a general pattern among animals.

Macronutrient consumption can also vary in response to environment and activity levels. For example, viral-infected caterpillars shifted toward a new self-selected macronutrient ratio that maximized survival (Cotter et al., 2011). During winter months (cold weather), golden snub-nosed monkeys increased their daily non-protein energy intake, probably due to the increased cost of thermoregulation (Guo et al., 2018). Many migratory birds adjust their nutrition to facilitate adequate fat accumulation (Bairlein, 2002). In early development, human energy requirements are highly correlated with basal metabolic rate and growth processes (0–6 month) (Butte et al., 2002). However, later when a child increases their physical activity, energy requirements correlate highly with activity level (reviewed in (Savarino et al., 2021)). For example, a single high intensity exercise increases lipid consumption in humans (Klausen et al., 1999).

Based on this literature, we hypothesized that animals will steadily reduce protein consumption during ontogeny because mass-specific growth rate declines (Brown et al., 2004; West et al., 2001; White et al., 2022), causing a progressive decrease in the consumption of protein relative to carbohydrate. We tested this hypothesis using South American locusts, *S. cancellata*. We predicted that mass-specific protein consumption would decrease strongly during development, in tight correlation with a decrease in mass-specific growth rate and a decrease in the protein:carbohydrate intake ratio, and that these relationships would hold across all animals because growth rate scales hypometrically across animals of different body sizes as well as during ontogeny (Hatton et al., 2019; Peters, 1983; West et al., 2001). We predicted that growth rates for field-collected animals placed on artificial diets would be similar to those of lab-reared animals, leading to similar protein consumption rates. However, because field-collected animals have experienced higher activity and stress levels, we predicted they would exhibit higher carbohydrate consumption and metabolic rates.

## Methods

### Locust lab culture

We used South American locusts (*Schistocerca cancellata*) from a captive colony at Arizona State University (ASU), 7–10 generations after locusts were collected from La Rioja and Catamarca regions of Argentina. The culture was kept at 30% RH, 34°C during the day and 25°C during the night, under 14 h light: 10 h dark photoperiod. Supplementary radiant heat was supplied during the daytime by incandescent 60 W electric bulbs next to the cages. In this general culture, locusts were fed daily with wheat shoots, fresh romaine lettuce leaves, and wheat bran *ad libitum*. For all experiments, Animals were excluded only if died during the experimental procedure.

### Artificial diets

The artificial diets were made as described by Dadd (Dadd, 1961) and adapted by Simpson and Abisgold (Simpson and Abisgold, 1985). We used five different isocaloric artificial foods in different assays that varied in protein and digestible carbohydrates: 7p:35c (% of protein and % of digestible carbohydrates, by dry mass), 14p:28c, 21p:21c, 28p:14c, 35p:7c. All the diets contained 54% cellulose and 4% vitamins and salts. The proteins were provided as a mix of 3:1:1 casein:peptone:albumen. The carbohydrate was provided as a 1:1 mix of sucrose and dextrin.

### Effect of ontogeny on intake targets

Nutritional intake targets were measured for each nymphal instar (1^st^–6^th^) (50 for each sex), from the first day of molting of each instar (or hatchlings for the 1^st^ instar nymphs) to the next molt. The adult intake (30 for each sex) targets were started on molt day and tested for three weeks. To have sufficient individuals of the same age, in each developmental stage, we monitored for newly molted individuals and randomly collected them on the same day. For the first instar nymphs, we monitored egg cups daily. When hatching was observed, we inserted the cups into standard colony rearing cages (45 × 45 × 45 cm metal mesh) for only a few hours to keep the ages of the nymphs as similar as possible. Sexing was performed by identifying presence/absence of developing ovipositor valves. For early developmental stages (1^st^–3^rd^ instars), we used a dissecting microscope to visualize these structures (SMZ-168, MOTIC, Schertz, TX, USA).

During these measurements, individuals were kept in plastic containers with holes drilled in the roof for ventilation which maintained the RH at ∼30%. The 1^st^ to 3^rd^ instar nymphs were kept in 11 × 16 × 4 cm cages, and 4^th^ instar nymphs to adults were kept in 19 × 10.4 × 14 cm containers. Each container had a water tube (refilled once a week), a perch (for successful molting) and two complementary artificial diets. To determine if locusts were arriving at a consistent p:c ratio and not just eating randomly from the two dishes, we provided half the locusts with the choice between 35p:7c and 7p:35c diets, while the other half were provided with the choice between 28p:14c and 7p:35c diets. We randomly placed cages from different diets pair treatments and sex on different shelves. To calculate consumption, we weighed each diet dish two times: 1) after drying and prior to inserting it into the assay boxes and 2) after it was removed from the experimental boxes and re-dried at 60°C for 24 h. To reduce error during the 1^st^ instar nymph experiment, we used small diet dishes (made from 1.5 ml Eppendorf lids) and weighed them with a micro balance (MSA6.6S-000-DM, accuracy of 10^-6^ g, Sartorius Weighing Technology GmbH, Goettingen, Germany). For all other instars, we used diet dishes made from an acrylic cylinder (10 × 25 mm) glued to a petri dish (58 mm in diameter) and weighed them using an analytical balance (accuracy of 10^-5^ g, XSE205, Mettler Toledo, Columbus, OH, US). Locusts were also weighed using the analytical balance. To reduce handling, which increases mortality, we weighed the locusts only after the experiment, and used final masses to correct consumption values. To calculate specific growth rate (equation 1) for each instar, we calculated mean initial masses for an extra 20 freshly molted (or newly hatched for first instar) individuals for each instar and sex.

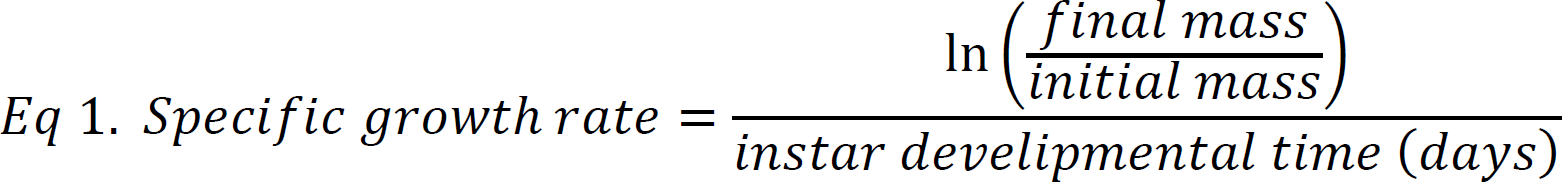

### Comparing intake targets and metabolic rates between lab-reared and field-captured locusts

We compared protein and carbohydrate consumption of 5^th^ and 6^th^ instar lab-reared nymphs (from the intake target experiment) to the field data we randomly collected from similarly aged nymphs in 2019 during the *S. cancellata* outbreak in Gran-Chaco, Paraguay (Talal et al., 2020). During the days of the experiments, field collected nymphs were kept at temperatures averaging 32.2 ± 1.94°C (measured with a Hobo logger, Onset, Bourne, MA, USA). We assessed macronutrient consumption rates by providing locusts with a choice between a low and a high carbohydrate diet (the same diets as in intake target experiment, see above) for eight days.

Comparison of resting metabolic rates was carried out on 6^th^ instar nymphs that were reared on confined artificial diets varying in protein:carbohydrate ratio (both in the field and in the lab) (Talal et al., 2021). Since we did not find an effect of dietary protein to carbohydrate ratio on oxygen consumption in either the lab or in the field (Talal et al., 2021), we pooled the data from the different diet treatment groups to compare resting metabolic rates between lab and field populations by measuring oxygen consumption. We carried out stop-flow respirometry using a FoxBox oxygen analyzer (Sable Systems International, Las Vegas, USA) as described in (Talal et al., 2021). Briefly, after inserting the nymph in a metabolic chamber and flushing it with CO_2_-free, dry, air, the chamber was sealed for period of time, after which a known volume was injected into CO_2-_free, dry air flow (500ml·min^-1^) which flushed through the oxygen analyzer. The metabolic rate (oxygen consumption) was temperature corrected to 34° C using Q10 of 2 (Talal et al., 2021).

### Statistics

Statistical analyses were performed using SPSS 20.0 (IBM) and R studio (Team, 2021). Prior to using parametric analyses, the normality of data was confirmed. For the intake target experiments: to rule out random feeding on different diet pairs, we employed multiple analysis of covariance (MANCOVA), using mass of carbohydrate and protein eaten as dependent variables, diet pair and sex as independent variables, and final body mass as a covariate. Due to some assumption violations, we compared protein and carbohydrate consumption rates as well as p:c ratios, among developmental stages and sexes, using aligned rank transformed observations on mass-specific values. To test for a significant effect of both developmental stage and sex we performed ANOVAs on aligned rank transformed observations according to the general procedure outlined by (Feys, 2016) using the software R (Team, 2021) and the R library ARTool (Matthew and Wobbrock, 2020).

To compare self-selected protein and carbohydrate consumption rates of lab-reared to field collected 5^th^ and 6^th^ instar nymphs, we used a Mann–Whitney *U* test (non-normal distribution). Oxygen consumption was measured in 6^th^ instar nymphs. We compared resting mass-specific oxygen consumption between field and lab 6^th^ instar nymphs using Mann–Whitney *U* tests (non-normal distribution).

## Results

### Protein to carbohydrate intake ratio decreased throughout development

We measured the self-selected protein:carbohydrate intake targets of each of the six nymphal stages and adults, for both males and females. Each individual was collected shortly after molting (or hatching from egg for first instar), sexed and relocated to an individual assay cage containing two pre-weighed diet dishes with chemically defined artificial diet varying in protein to carbohydrate ratios. We determined self-selected macronutrient consumption and ratios by reweighing the diet dishes following individual molting to the next developmental stage (or following three weeks of maturation in adults) (Fig. 1). We found that younger instars (1^st^ to 4^th^) had protein-biased consumption (selected high p:c, protein to carbohydrate ratios) with 3^rd^ instar nymphs exhibiting the highest p:c of 1.37p:1c (Fig. 1D). In contrast, older locusts became carbohydrate-biased, and adults selected 1p:2.66c intake targets (Fig. 1D). Males and females (both unmated) did not differ from each other in relative macronutrient consumption during most of the nymphal stages (Fig. 1 and Table 1). There were no significant interactions between sex and diet pairs on total macronutrient consumption (Table 1).

**Figure 1.**
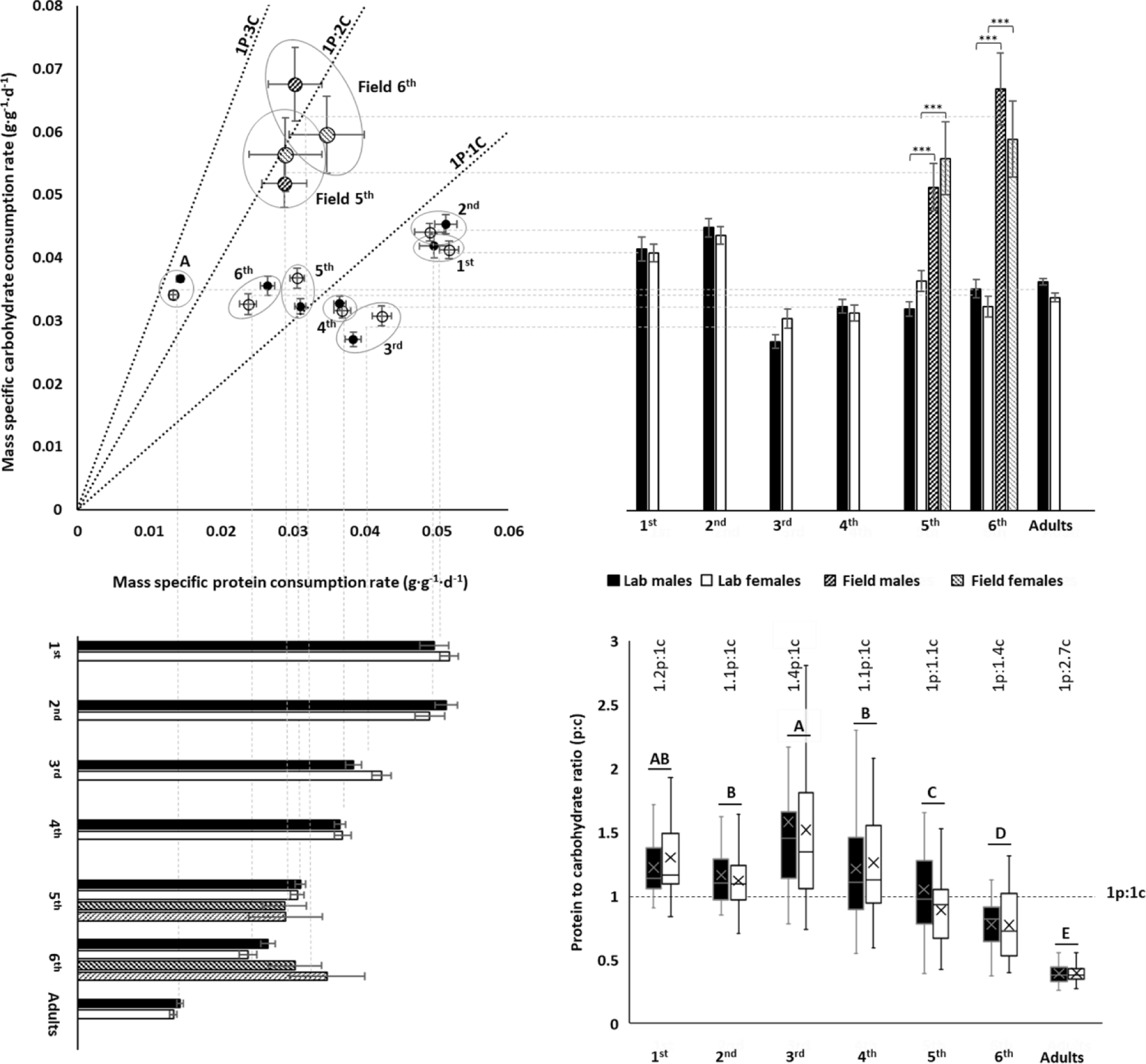
**A**. Self-selected protein to carbohydrate (p:c) consumption rates decreased systematically during ontogeny. **B**. For lab-reared locusts, mass-specific carbohydrate consumption rates were highest in early instars relative to older instars and adults. Field-collected 5^th^ and 6^th^ instars consumed more carbohydrate than lab-reared nymphs. **C**. For lab-reared locusts, mass-specific protein consumption declined systematically with age. Field-collected 5^th^ and 6^th^ instar nymphs consumed protein at similar rates to lab-reared animals. **D**. Young, 1^st^ – 4^th^ nymph instars self-selected protein biased intake target ratios, whereas later in development, locusts became carbohydrate biased (medians and interquartile ranges are represented by the boxes and center line, with an X to indicate the mean). The numbers above the boxes represent life stage averaged (both sexes) p:c intake targets. The posthoc letters were given only when there was not significant interactive developmental stage * sex effect. For panels A, B, and C, means and standard errors (SEM) are shown. The three asterisks represent significant differences between lab and field populations when p < 0.001. Throughout, males are black circles/bars and females are white circles/bars; field locusts are represented by striped bars.

**Table 1.** The diet pair presented did not affect the amount of protein and carbohydrate consumed at any developmental stage (multiple analysis of covariance (MANCOVA), with diet pairs as blocks and masses as a covariate).

| Nymph instar | | F-value | p-value | Wilks' $\Lambda$ |
| --- | --- | --- | --- | --- |
| 1<sup>st</sup> | Diet | F(2,72)=2.82 | 0.07 | 0.93 |
|  | Sex | F(2,72)=1.46 | 0.24 | 0.96 |
|  | Diet X Sex | F(2,72)=0.36 | 0.7 | 0.99 |
| 2<sup>nd</sup> | Diet | F(2,91)=2.33 | 0.1 | 0.95 |
|  | Sex | F(2,91)=2.63 | 0.08 | 0.95 |
|  | Diet X Sex | F(2,91)=0.10 | 0.91 | 0.99 |
| 3<sup>rd</sup> | Diet | F(2,93)=1.09 | 0.34 | 0.98 |
|  | Sex | F(2,93)=10.1 | <0.001 | 0.82 |
|  | Diet X Sex | F(2,93)=0.87 | 0.42 | 0.98 |
| 4<sup>th</sup> | Diet | F(2,91)=0.38 | 0.68 | 0.99 |
|  | Sex | F(2,91)=2.84 | 0.06 | 0.94 |
|  | Diet X Sex | F(2,91)=0.32 | 0.73 | 0.99 |
| 5<sup>th</sup> | Diet | F(2,84)=1.17 | 0.32 | 0.97 |
|  | Sex | F(2,84)=2.84 | 0.06 | 0.94 |
|  | Diet X Sex | F(2,84)=0.82 | 0.44 | 0.98 |
| 6<sup>th</sup> | Diet | F(2,48)=0.91 | 0.41 | 0.96 |
|  | Sex | F(2,48)=1.18 | 0.32 | 0.95 |
|  | Diet X Sex | F(2,48)=1.37 | 0.26 | 0.95 |
| Adult | Diet | F(2,47)=0.752 | 0.477 | 0.969 |
|  | Sex | F(2,47)=5.191 | 0.009 | 0.819 |
|  | Diet X Sex | F(2,47)=0.184 | 0.832 | 0.992 |

Mass-specific carbohydrate consumption rates were about 30% higher for the first two instars compared to older animals (ANOVA: diet: F_6,554_ = 34.459; p < 0.001, Fig. 1B). Males and females did not differ significantly in mass-specific carbohydrate consumption rates (ANOVA: sex: F_1,554_ = 0.294; p = 0.940, Fig. 1B). Mass-specific protein consumption rate decreased steadily through ontogeny, with a roughly four-fold decrease in adults compared to first instars (ANOVA: diet: F_6,554_ = 193.142; p < 0.001, Fig. 1C). There were differences between the sexes (ANOVA: sex: F_1,554_ = 7.055; p = 0.008) and a significant interactive sex * diet effect on mass-specific protein consumption (ANOVA: sex*diet: F_6,554_ = 38.995; p = 0.011) which was associated with small, irregular stage-effects on which sex consumed more. Together, these ontogenetic effects on carbohydrate and protein consumption led to strong decreases in the protein:carbohydrate intake ratio through ontogeny, with the youngest instars consuming about 30% more protein than carbohydrate and the oldest juveniles and adults consuming approximately twice as much carbohydrate as protein (ANOVA: Sex: F_1,574_ = 3.112, p = 0.078;Developmental stage: F_6,574_ = 87.529, p < 0.001; Sex* Developmental stage: F_6,574_ = 1.419, p = 0.645) (Fig. 1D).

### Protein, but carbohydrate consumption correlates with growth and scales hypometrically

The decrease in protein consumption and requirements during development could be explained by the decrease in specific growth rates, which were highly correlated with mass-specific protein consumption in both sexes (Fig. 2A). Conversely, mass-specific carbohydrate consumption was uncorrelated with specific growth rates (Fig. 2B). Plotting macronutrient consumption rates on a log-log plot revealed strong correlation with body mass (Fig. 3). Whereas protein consumption rates in *S. cancellata* scaled strongly hypometrically, with slope of 0.761 (95% confidence interval: 0.744–0.778) (Fig. 3A), carbohydrate consumption rates scaled hypometrically, but with a slope near 1 (slope of 0.939; 95% confidence interval: 0.92–0.957) (Fig. 3B).

**Figure 2.**
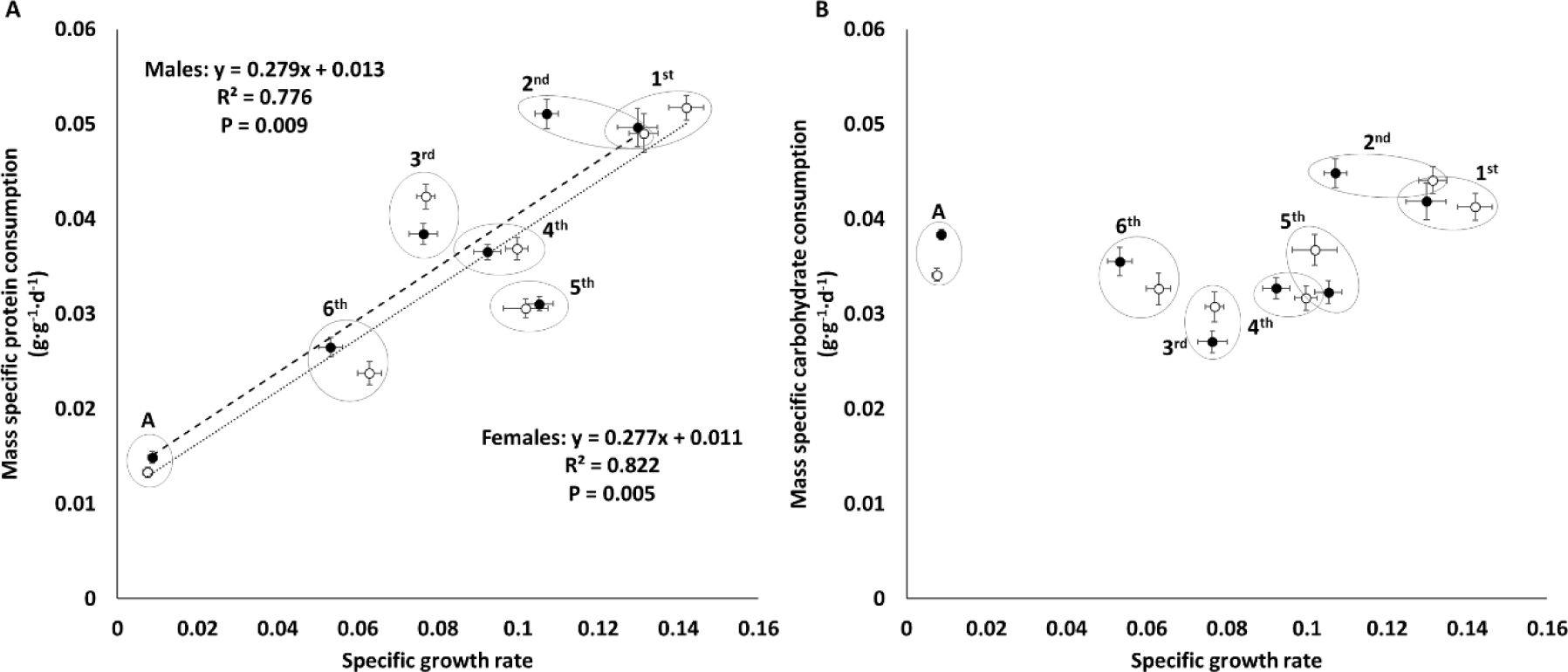
Mass-specific protein consumption rate was well-predicted by specific growth rate across ontogeny (**A**), whereas mass-specific carbohydrate consumption rate was not (**B**). Filled circles and dashed line represent males, whereas opened circles and dotted line represent females. Means and standard errors (SEM) are shown.

**Figure 3.**
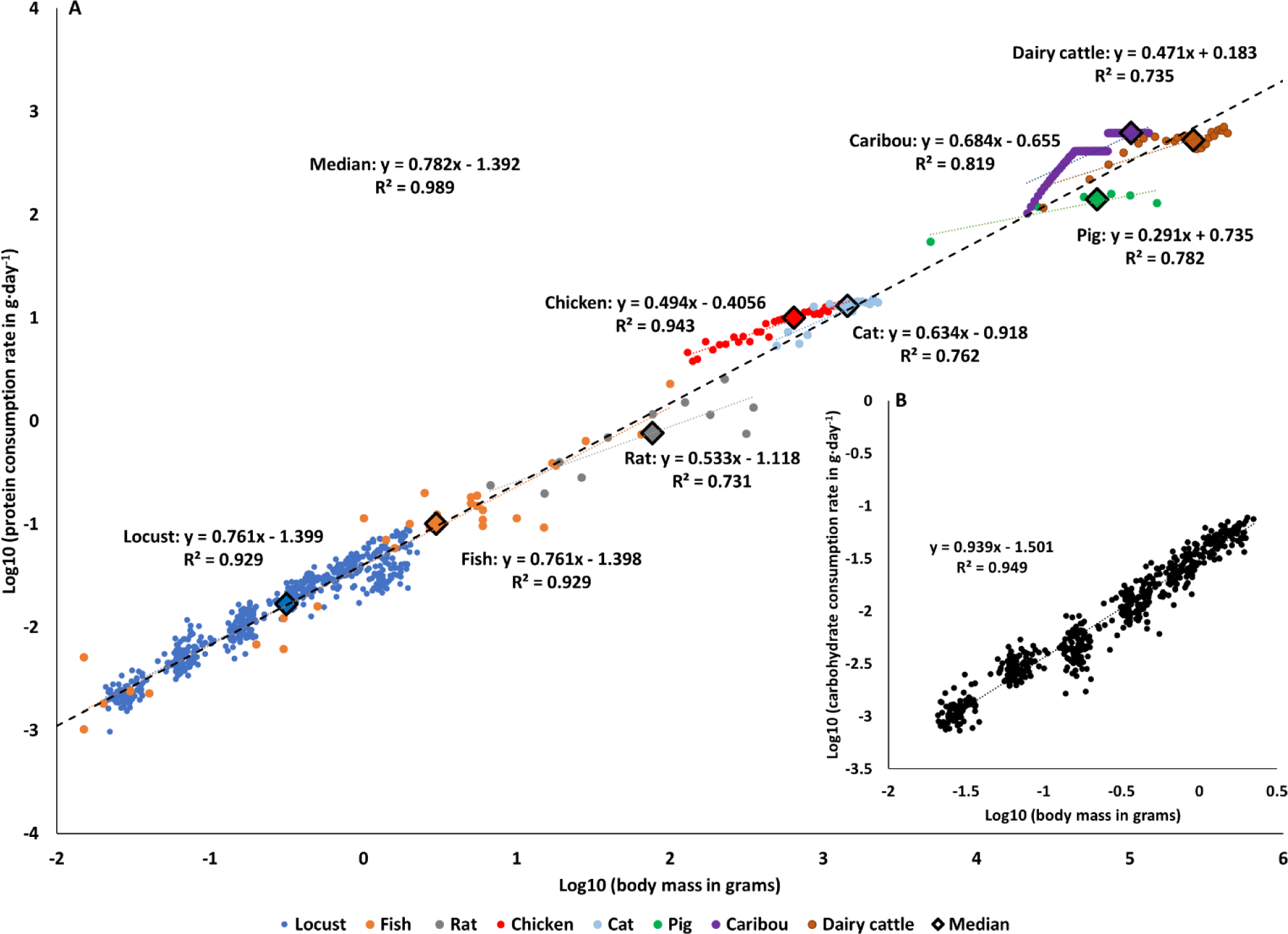
**A.** Protein consumption rate scales hypometrically throughout development across the animal kingdom. The blue circles: locusts (*Schistocerca cancellata*, this study); orange circles: fish (early development in multiple species (reviewed in (Dabrowski, 1986)); gray circles: rats (*Rattus rattus* (Ricci and Levin, 2003)); red circles: chicken (*Gallus gallus domesticus* (Kaufman et al., 1978)); light blue circles: cats (*Felis catus* (Dickinson and Scott, 1956; Miller and Allison, 1958)); green circles: pigs (*Sus domesticus* (Black et al., 1986)); purple circles: caribou (*Rangifer tarandus* (McEwan, 1968)); brown circles: dairy cattle (*Bos taurus* (Crichton et al., 1959)). Diamonds represent the median value of each taxonomic group (matched by color), and the black dashed line is the across species regression. The data were corrected to 37°C using a Q10 of 2. **B.** Carbohydrate consumption rates scales hypometrically, a very close to isometrically in *Schistocerca cancellata*.

### Field collected nymphs had higher rates of metabolism and carbohydrate consumption but similar protein consumption as lab-reared locusts

Field-collected South American locusts had more carbohydrate-biased intake targets relative to lab-reared locusts (Fig 1A). Male 5^th^ and 6^th^ instar nymphs collected from field populations (Gran Chaco, Paraguay, April 2019) had 50–90% higher carbohydrate consumption rates relative to lab-reared nymphs (Mann–Whitney *U* test: *U* = 2; *U* = 17; respectively; p < 0.001 for both instars) as did female 5^th^ and 6^th^ instar nymphs (Mann–Whitney *U* test: *U* = 6; *U* = 2; respectively; p < 0.001 for both instars) (Fig. 1A and 1B). However, there were no significant differences in protein consumption between field collected and lab reared nymphs, for male 5^th^ and 6^th^ instar nymphs (Mann–Whitney *U* test: *U* = 204; p = 0.197; *U* = 163; p = 0.135; respectively) or female 5^th^ and 6^th^ instar nymphs (Mann–Whitney *U* test: *U* = 43; p = 0.071; *U* = 127; p = 0.859; respectively) (Fig. 1B and 1C). The higher carbohydrate consumption of field-captured locusts was partly due to a higher resting metabolic rate. Using stop-flow respirometry we demonstrated that field collected 6^th^ (N = 29) instar nymphs had ∼23% higher mass-specific resting oxygen consumption rate than 6^th^ (N = 50) instar lab-reared nymphs (1.126 ± 0.052 ml·g^-1^·h^-1^, 0.914 ± 0.021 ml·g^-1^·h^-1^, mean ± SEM for field-collected and lab-reared, respectively) (Mann–Whitney *U* test: *U* = 349; p < 0.001).

## Discussion

Overall, our results demonstrate that macronutrient targets change predictably from high protein:carbohydrate consumption in the young toward increasingly lower protein: carbohydrate intake targets during ontogeny in *S. cancellata*. From first instar to adult *S. cancellata*, mass-specific protein consumption rate decreased fourfold with little change in mass-specific carbohydrate consumption (Fig. 1). The decrease in mass-specific protein consumption rate was tightly correlated with a decline in specific growth rate, likely explaining the shift in protein requirements (Fig. 2A). In contrast, protein demand did not differ between lab and field populations, but carbohydrate consumption rate was >50% higher in field populations (Fig. 1).

A decline in protein consumption during ontogeny could be a common, and likely a general pattern for animals. Combining our data with the available literature for animals revealed that declining protein consumption rates during ontogeny or across species is a general pattern for animals (Fig. 3A). Older and larger animals consume proportionally less protein in locusts, fish, rats, chickens, pigs, cats, caribou, and dairy cattle, and with a very similar pattern holding across species (Fig. 3A). Thus, protein consumption can be added to the list of traits that scale predictably with body size (Schmidt-Nielsen, 1995; Sibly et al., 2012). The hypometric scaling of protein consumption across species is consistent with the general hypometric scaling of growth rates across animals (Hatton et al., 2019). Though ontogenetic slopes of protein consumption on mass were much lower than the cross-species pattern in a few groups, including cats and pigs; it will be interesting to determine whether such variation relates to interspecific variation in the scaling of ontogenetic growth and lifespan.

Assuming that energy needs and consumption are primarily set by metabolic rate, we would expect that mass specific non-protein consumption to decrease with both animal mass and age due to the generally-observed hypometric scaling of metabolic rates across animal taxa (Harrison et al., 2022; White et al., 2022). In locusts, we demonstrated that carbohydrate consumption scaled hypometrically, but with a slope very close to 1, a much higher mass-scaling exponent than observed for protein consumption (Fig. 3A), but in the range of reported scaling for resting metabolic rate (0.77-1) for orthopterans (Fielding and DeFoliart, 2008; Greenlee and Harrison, 2004). Likely, in locusts, carbohydrate consumption of older individuals is increased due to the increase in mass-specific lipid stores that occurs in older juveniles and adults, as stored lipids are mainly synthesized from ingested carbohydrates (Talal et al., 2021).

An important goal for the field of nutritional ecology is to predict nutritional needs, foraging behavior and strategies, and consequences of nutritional imbalance for animals in the field (Behmer and Joern, 2008). Relative to the lab population, we measured a 50-90% increase in carbohydrate consumption rates for field-collected 5^th^ and 6^th^ instar nymphs. In contrast, protein consumption rates did not vary between lab and field in our study. This may not be true under every ecological condition; for example, poor resource conditions that reduce growth will likely also reduce protein consumption. Nonetheless, these data support the hypothesis that protein-consumption rates of animals in good field conditions may be predicted from results with lab-reared animals. Captive animals usually do not need to actively forage, which can be energetically expensive and cause long-term effects on resting metabolic rates (Bergman et al., 2001). Studies of monkeys and apes have demonstrated that decreases in foraging activity in captivity may promote metabolic suppression, diabetes, and obesity (reviewed in (Bellisari, 2008)). Increased energy demands and energy metabolism in field animals may also be due to a past history of consumption of tougher, better chemically defended plants (Clissold et al., 2009; Maskato et al., 2014).

## Conclusions and Future Directions

Hypometric scaling of protein consumption is associated with declining specific growth rate during ontogeny and across species in animals, providing a new and useful paradigm for nutritional ecology. Many important questions remain. Is species-level variation in the ontogenetic scaling of protein consumption rate predictable by species differences in growth rates? How do ecological conditions that affect growth and reproduction affect this pattern? How useful would age-specific diets be for humans and animal husbandry? How will warming climate and increases in body temperatures of ectotherms affect the nutritional needs of animals, and their interactions with natural plants and microbial communities?

## Acknowledgments

We thank Kelly O’Meara and Geoffrey Osgood, our former lab technicians, for helping with organization and logistics. Aunmolpreet Chahal assisting with experimental setup, as well as providing the locusts with diets and water tubes. Special thanks for Mai and Tom Talal for cleaning hundreds of experimental cages, water tubes, and diet dishes, when experiments were finished. We also thank out South American collaborators for helping with collecting the colony and helping with field logistics: Eduardo V. Trumper (INTA, Argentina), Luis Sanchez Shimura (SENASAG, Bolivia), Fernando Copa Bazán (Universidad Autónoma Gabriel René Moreno, Bolivia), Jorge Frana (INTA, Argentina) and Julio E. Rojas (SENAVE, Paraguay). In the US, the authors recognize that the ASU campus community has and continues to benefit from land that was taken from Indigenous communities, including the Akimel O’odham (Pima) and Pee Posh (Maricopa) Indian Communities, whose stewardship of these lands allows us to be here today.

## Author Contributions

S.T.: conceptualization, data curation, formal analysis, investigation, methodology, visualization, writing original draft and reviewing and editing. J.F.H.: conceptualization, funding acquisition, investigation, methodology, project administration, resources, supervision, writing-reviewing and editing. R.F.: investigation, methodology, writing original draft and reviewing and editing. J.P.Y.: investigation, methodology, writing original draft and reviewing and editing. H.E.M.: investigation, methodology, writing original draft and reviewing and editing. R.O.: investigation, methodology, writing original draft and reviewing and editing. A.J.C.: conceptualization, funding acquisition, investigation, methodology, project administration, resources, supervision, writing-reviewing and editing.

## Figure 1 – Source data file 1

This source file contains separate panels for figure 1, including sample sizes and states. The raw data for these figure panels, as well as for the rest of the figures and other data, is attached as supplementary material file.

**Panel A:**
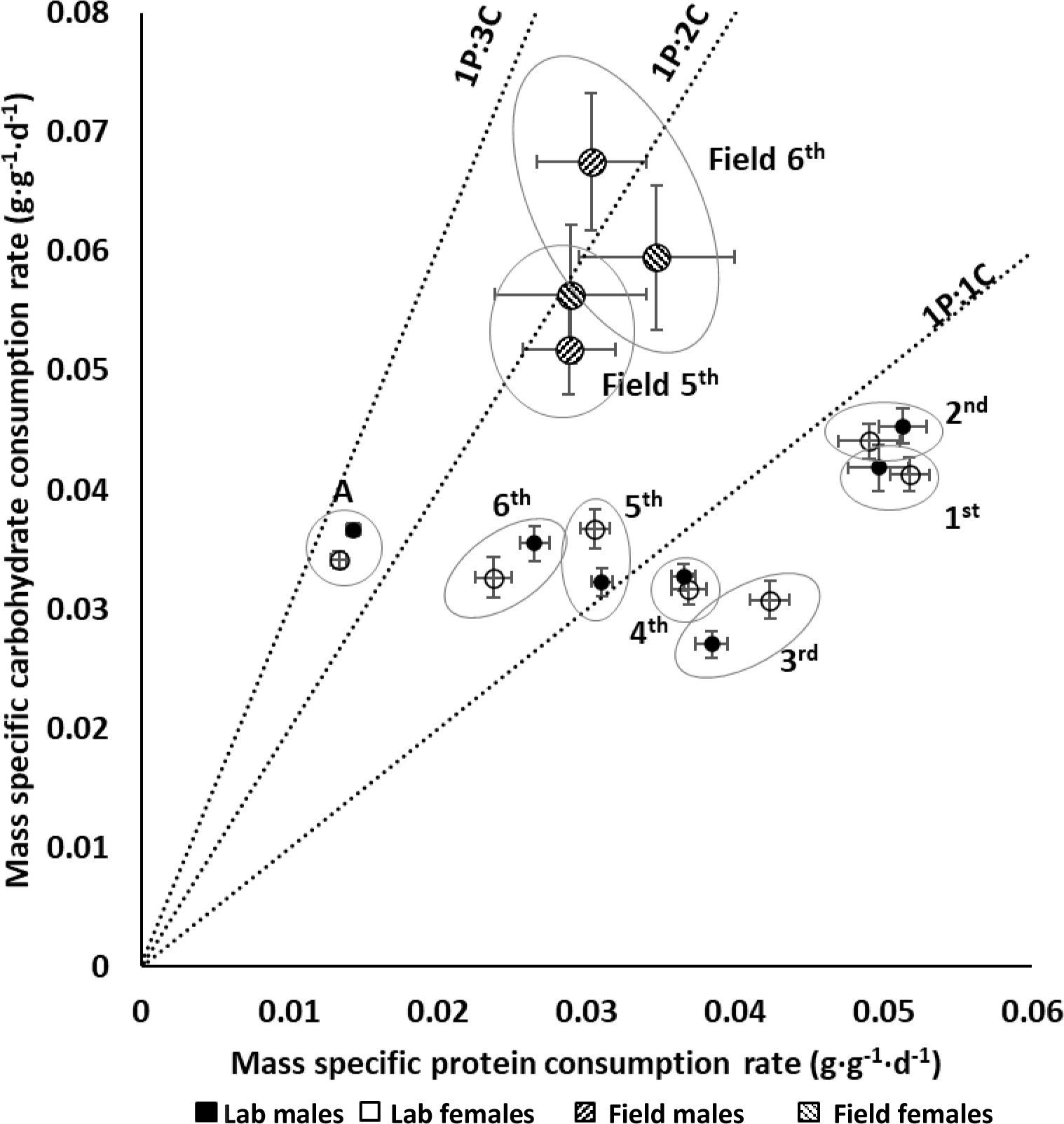
Self-selected protein to carbohydrate (p:c) consumption rates decreased systematically during ontogeny (For sample sizes see supplementary table 1, at the end of this document).

**Panel B:**
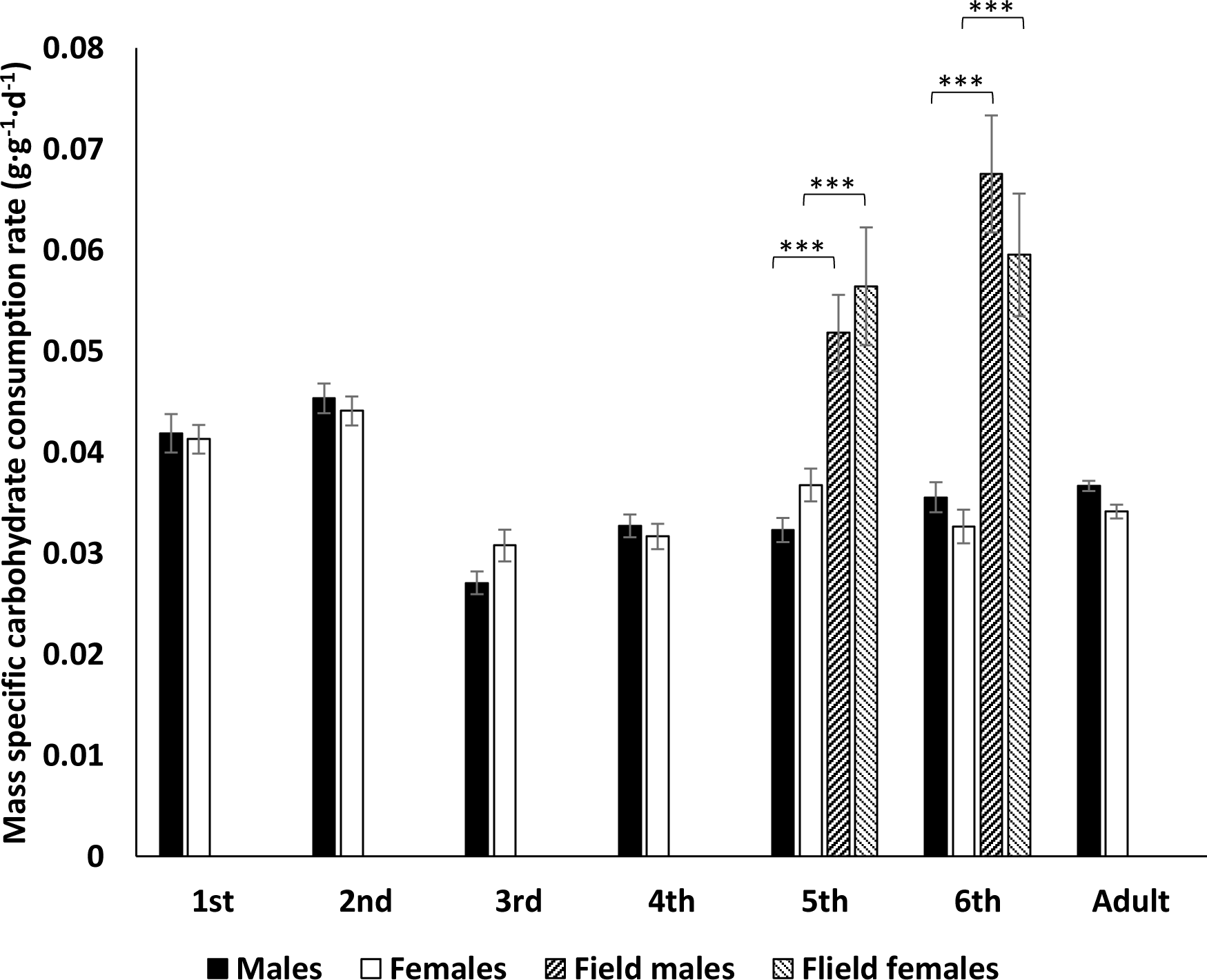
Lab males and females did not differ significantly in mass-specific carbohydrate consumption rates (ANOVA: sex: F_1,554_ = 0.294; p = 0.940, Fig. 1B).Mass-specific carbohydrate consumption rates were significantly different among different developmental stages (ANOVA: diet: F_6,554_ = 34.459; p < 0.001, Fig. 1B) and there was significant interactive sex * diet effect on mass-specific carbohydrate consumption (ANOVA: sex*diet: F_6,554_ = 2.363; p = 0.029). Male 5^th^ and 6^th^ instar nymphs collected from field populations (Gran Chaco, Paraguay, April 2019) had 50–90% higher carbohydrate consumption rates relative to lab-reared nymphs (Mann–Whitney U test: U = 2; U = 17; respectively; p < 0.001 for both instars) as did female 5^th^ and 6^th^ instar nymphs (Mann–Whitney U test: U = 6; U = 2; respectively; p < 0.001 for both instars).

**Panel C:**
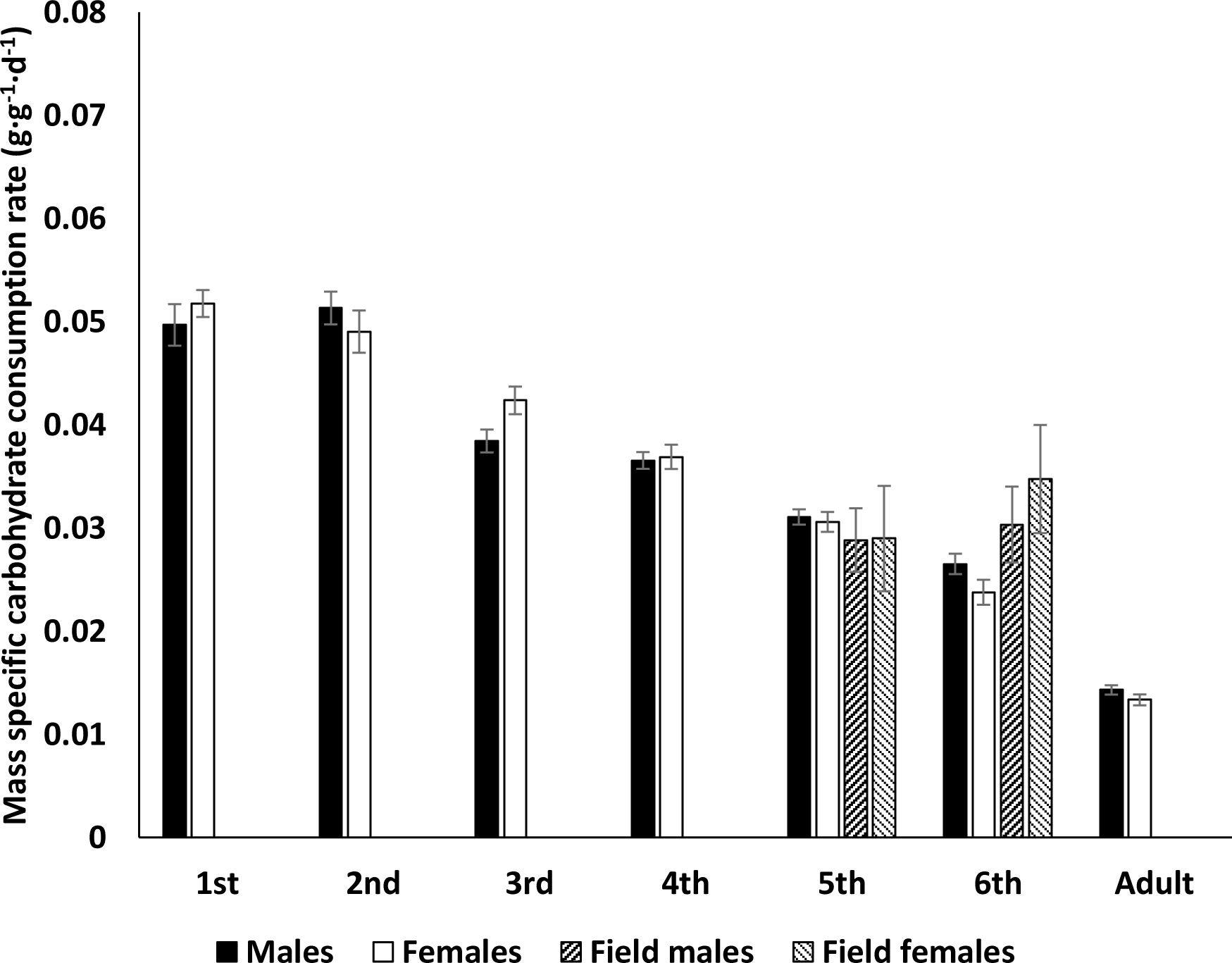
In lab population mass-specific protein consumption rate decreased steadily through ontogeny (ANOVA: diet: F_6,554_ = 193.142; p < 0.001, Fig. 1C). There were differences between the sexes (ANOVA: sex: F_1,554_ = 7.055; p = 0.008) and a significant interactive sex * diet effect on mass-specific protein consumption (ANOVA: sex*diet: F_6,554_ = 38.995; p = 0.011). There were no significant differences in protein consumption between field collected and lab reared nymphs, for male 5^th^ and 6^th^ instar nymphs (Mann–Whitney U test: U = 204; p = 0.197; U = 163; p = 0.135; respectively) or female 5^th^ and 6^th^ instar nymphs (Mann–Whitney U test: U = 43; p = 0.071; U = 127; p = 0.859; respectively).

**Panel D:**
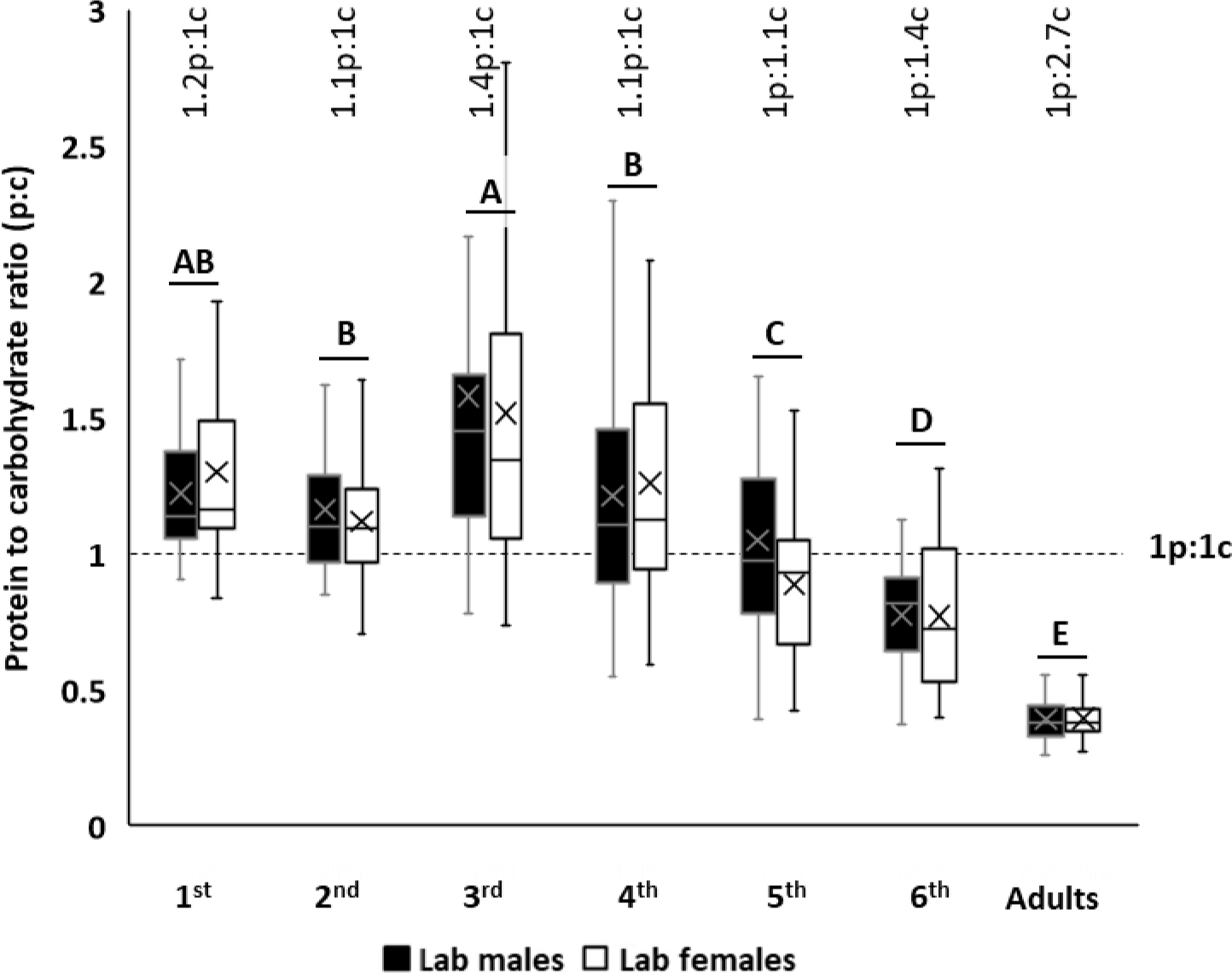
There were strong decreases in the protein:carbohydrate intake ratio through ontogeny (ANOVA: Sex: F1,574 = 3.112, p = 0.078; Developmental stage: F6,574 = 87.529, p < 0.001; Sex* Developmental stage: F6,574 = 1.419, p = 0.645).

**Supplementary table 1.** Sample sizes of the lab and field populations.

| **Lab population** |  |  |
| --- | --- | --- |
| **Developmental stage** | **Sex** | **Sample size** |
| 1<sup>st</sup> instar nymph | Male | 37 |
|  | Female | 41 |
| 2<sup>nd</sup> instar nymph | Male | 46 |
|  | Female | 51 |
| 3<sup>rd</sup> instar nymph | Male | 50 |
|  | Female | 48 |
| 4<sup>th</sup> instar nymph | Male | 58 |
|  | Female | 40 |
| 5<sup>th</sup> instar nymph | Male | 55 |
|  | Female | 35 |
| 6<sup>th</sup> instar nymph | Male | 30 |
|  | Female | 24 |
| Adult | Male | 28 |
|  | Female | 25 |

| **Field population** |  |  |
| --- | --- | --- |
| **Developmental stage** | **Sex** | **Sample size** |
| 5<sup>th</sup> instar nymph | Male | 10 |
|  | Female | 15 |
| 6<sup>th</sup> instar nymph | Male | 5 |
|  | Female | 11 |

## References

Babaei SS, Abedian Kenari A, Nazari R, Gisbert E. 2011. Developmental changes of digestive enzymes in Persian sturgeon (Acipenser persicus) during larval ontogeny. Aquaculture 318:138–144. doi:10.1016/j.aquaculture.2011.04.032

Bairlein F. 2002. How to get fat: nutritional mechanisms of seasonal fat accumulation in migratory songbirds. Naturwissenschaften 89:1–10. doi:10.1007/s00114-001-0279-6

Ballard O, Morrow AL. 2013. Human milk composition, nutrition and bioactive factors. Pediatr Clin NA 60:49–74. doi:10.1016/j.pcl.2012.10.002

Behmer S, Joern A. 2008. Coexisting generalist herbivores occupy unique nutritional feeding niches. Proc Natl Acad Sci 105:1977–1982. doi:10.1073/pnas.0711870105

Behmer ST. 2009. Insect herbivore nutrient regulation. Annu Rev Entomol 54:165–187. doi:doi:10.1146/annurev.ento.54.110807.090537

Bellisari A. 2008. Evolutionary origins of obesity. Obes Rev 9:165–180. doi:https://doi.org/10.1111/j.1467-789X.2007.00392.x

Bergman CM, Fryxell JM, Gates CC, Fortin D. 2001. Ungulate foraging strategies: energy maximizing or time minimizing? J Anim Ecol 70:289–300. doi:https://doi.org/10.1111/j.1365-2656.2001.00496.x

Black JL, Campbell RG, Williams IH, James K., Davies GT. 1986. Simulation of energy and amino acid utilisation in the pig. Res Dev Agric 3:121–145.

Bouchard SS, Bjorndal KA. 2006. Ontogenetic diet shifts and digestive constraints in the omnivorous freshwater turtle Trachemys scripta. Physiol Biochem Zool 79:150–158. doi:10.1086/498190

Brown JH, Gillooly JF, Allen AP, Savage VM, West GB. 2004. Toward a metabolic theory of ecology. Ecology 85:1771–1789.

Butte NF, Mardia G. Lopez-Alarcon, Cutberto G. 2002. Nutrient adequacy of exclusive breastfeeding for the term infant during the first six months of life. World Health Organization.

Cease AJ, Elser JJ, Ford CF, Hao S, Kang L, Harrison JF. 2012. Heavy livestock grazing promotes locust outbreaks by lowering plant nitrogen. Science (80- ) 335:467–469.

Clissold FJ, Sanson GD, Read J, Simpson SJ. 2009. Gross vs. net income: How plant toughness affects performance of an insect herbivore. Ecology 90:3393–3405.

Cohen RW, Heydon SL, Waldbauer GP, Friedman S. 1987. Nutrient self-selection by the omnivorous cockroach Supella longipalpa. J Insect Physiol 33:77–82. doi:https://doi.org/10.1016/0022-1910(87)90077-1

Cotter SC, Simpson SJ, Raubenheimer D, Wilson K. 2011. Macronutrient balance mediates trade-offs between immune function and life history traits. Funct Ecol 25:186–198.

Crichton JA, Aitken JN, Boyne AW. 1959. The effect of plane of nutrition during rearing on growth, production, reproduction and health of dairy cattle. I. Growth to 24 months. Anim Sci 1:145–162. doi:DOI: 10.1017/S0003356100033353

Dabrowski KR. 1986. Ontogenetical aspects of nutritional requirements in fish. Comp Biochem Physiol Part A Physiol 85:639–655. doi:https://doi.org/10.1016/0300-9629(86)90272-0

Dadd RH. 1961. The nutritional requirements of locusts-IV. Requirements for vitamins of the B complex. J Ins Physiol 6:1–12.

Dickinson CD, Scott PP. 1956. Nutrition of the cat: 1. A practical stock diet supporting growth and reproduction. Br J Nutr 10:304–311. doi:DOI: 10.1079/BJN19560046

Feys J. 2016. Nonparametric tests for the interaction in two-way factorial designs using R. R J 367–378.

Fielding DJ, DeFoliart LS. 2008. Relationship of metabolic rate to body size in Orthoptera. J Orthoptera Res 17:301–306.

Fink P, Pflitsch C, Marin K. 2011. Dietary essential amino acids affect the reproduction of the keystone herbivore Daphnia pulex. PLoS One 6:e28498.

Ghabrial A, Luschnig S, Metzastein M, Krasnov m. 2011. Longitudinal analysis of macronutrients and minerals in human milk produced by mothers of preterm infants. Clin Nutr 30:215–220. doi:10.1016/j.clnu.2010.08.003

Greenlee KJ, Harrison JF. 2004. Development of respiratory function in the American locust Schistocerca americana I. Across-instar effects. J Exp Biol 207:497–508.

Guo S-T, Hou R, Garber PA, Raubenheimer D, Righini N, Ji W-H, Jay O, He S-J, Wu F, Li F-F, Li B-G. 2018. Nutrient-specific compensation for seasonal cold stress in a free-ranging temperate colobine monkey. Funct Ecol 32:2170–2180. doi:https://doi.org/10.1111/1365-2435.13134

Harrison JF, Biewener A, Bernhardt JR, Burger JR, Brown JH, Coto ZN, Duell ME, Lynch M, Moffett ER, Norin T, Pettersen AK, Smith FA, Somjee U, Traniello JFA, Williams TM. 2022. White Paper: An Integrated Perspective on the Causes of Hypometric Metabolic Scaling in Animals. Integr Comp Biol icac136. doi:10.1093/icb/icac136

Hatton IA, Dobson AP, Storch D, Galbraith ED, Loreau M. 2019. Linking scaling laws across eukaryotes. Proc Natl Acad Sci 116:21616–21622. doi:10.1073/pnas.1900492116

Hewson-Hughes AK, Hewson-Hughes VL, Colyer A, Miller AT, McGrane SJ, Hall SR, Butterwick RF, Simpson SJ, Raubenheimer D. 2013. Geometric analysis of macronutrient selection in breeds of the domestic dog, Canis lupus familiaris. Behav Ecol 24:293–304. doi:10.1093/beheco/ars168

Hewson-Hughes AK, Hewson-Hughes VL, Miller AT, Hall SR, Simpson SJ, Raubenheimer D. 2011. Geometric analysis of macronutrient selection in the adult domestic cat, felis catus. J Exp Biol 214:1039–1041. doi:10.1242/jeb.049429

Karameta E, Mizan VL, Sagonas K, Sfenthourakis S, Valakos ED, Pafilis P. 2017. Ontogenetic shifts in the digestive efficiency of an insectivorous lizard (Squamata: Agamidae). Salamandra 53:321–326.

Kaufman LW, Collier G, Squibb RL. 1978. Selection of an adequate protein-carbohydrate ratio by the domestic chick. Physiol Behav 20:339–344. doi:https://doi.org/10.1016/0031-9384(78)90229-9

Klausen B, Toubro S, Ranneries C, Rehfeld JF, Holst JJ, Christensen NJ, Astrup A. 1999. Increased intensity of a single exercise bout stimulates subsequent fat intake. Int J Obes 23:1282–1287. doi:10.1038/sj.ijo.0801074

Knoff A, Hohn A, Macko S. 2008. Ontogenetic diet changes in bottlenose dolphins (Tursiops truncatus) reflected through stable isotopes. Mar Mammal Sci 24:128–137. doi:https://doi.org/10.1111/j.1748-7692.2007.00174.x

Lee KP, Simpson SJ, Raubenheimer D. 2004. A comparison of nutrient regulation between solitarious and gregarious phases of the specialist caterpillar, Spodoptera exempta (Walker). J Insect Physiol 50:1171–1180. doi:https://doi.org/10.1016/j.jinsphys.2004.10.009

Maskato Y, Talal S, Keasar T, Gefen E. 2014. Red foliage color reliably indicates low host quality and increased metabolic load for development of an herbivorous insect. Arthropod Plant Interact 8:285–292. doi:10.1007/s11829-014-9307-2

Matthew K, Wobbrock O. 2020. Aligned Rank Transform for Nonparametric Factorial ANOVAs. Conf Hum Factors Comput Syst 143–146.

McEwan EH. 1968. Growth and development of the barren-ground caribou. II. Postnatal growth rates. Can J Zool 46:1023–1029.

Miller SA, Allison JB. 1958. The Dietary Nitrogen Requirements of the Cat. J Nutr 64:493–501. doi:10.1093/jn/64.3.493

Nicholas K, Simpson K, Wilson M, Trott J, Shaw D. 1997. The Tammar Wallaby: A Model to Study Putative Autocrine-Induced Changes in Milk Composition. J Mammary Gland Biol Neoplasia 2:299–310. doi:10.1023/A:1026392623090

Peters RH. 1983. The Ecological Implications of Body Size. Cambridge: Cambridge University Press.

Raubenheimer D, Senior AM, Mirth C, Cui Z, Hou R, Le Couteur DG, Solon-Biet SM, Léopold P, Simpson SJ. 2022. An integrative approach to dietary balance across the life course. iScience 25:104315. doi:https://doi.org/10.1016/j.isci.2022.104315

Ricci MR, Levin BE. 2003. Ontogeny of diet-induced obesity in selectively bred Sprague-Dawley rats. Am J Physiol Integr Comp Physiol 285:R610–R618. doi:10.1152/ajpregu.00235.2003

Riedman M, Ortiz CL. 1979. Changes in Milk Composition during Lactation in the Northern Elephant Seal. Physiol Zool 52:240–249. doi:10.1086/physzool.52.2.30152567

Savarino G, Corsello A, Corsello G. 2021. Macronutrient balance and micronutrient amounts through growth and development. Ital J Pediatr 47:109. doi:10.1186/s13052-021-01061-0

Schmidt-Nielsen K. 1995. Scaling: Why is Animal Size So Important? Cambridge: Cambridge University Press.

Sibly RM, Brown JH, Kodric-Brown A. 2012. Metabolic Ecology: A Scaling Approach. Wiley-Blackwell.

Simpson SJ, Abisgold JD. 1985. Compensation by locusts for changes in dietary nutrients: behavioural mechanisms. Physiol Entomol 10:443–452.

Simpson SJ, Raubenheimer D. 2012. The nature of nutrition. A unifying framework from animal adaptation to human obisity. Princeton University Press.

Stockhoff BA. 1993. Ontogenetic change in dietary selection for protein and lipid by gypsy moth larvae. J Insect Physiol 39:677–686. doi:10.1016/0022-1910(93)90073-Z

Talal S, Cease A, Farington R, Medina HE, Rojas J, Harrison J. 2021. High carbohydrate diet ingestion increases post-meal lipid synthesis and drives respiratory exchange ratios above 1. J Exp Biol 224:jeb240010. doi:10.1242/jeb.240010

Talal S, Cease AJ, Youngblood JP, Farington R, Trumper E V, Medina HE, Rojas JE, Fernando Copa A, Harrison JF. 2020. Plant carbohydrate content limits performance and lipid accumulation of an outbreaking herbivore. Proc R Soc B Biol Sci 287:20202500. doi:10.1098/rspb.2020.2500

Team RC. 2021. R: A language and environment for statistical computing. R Found Stat Comput Vienna, Austria.

West GB, Brown JH, Enquist BJ. 2001. A general model for ontogenetic growth. Nature 413:628–631.

White CR, Alton LA, Bywater CL, Lombardi EJ, Marshall DJ. 2022. Metabolic scaling is the product of life-history optimization. Science (80-) 377:834–839. doi:10.1126/science.abm7649

